# Cyclic Electromagnetic DNA Simulation (CEDS) influences the functions of Long Oligo-dsDNAs and Plasmid DNAs

**DOI:** 10.1101/2024.08.24.609489

**Authors:** Suk Keun Lee, Dae Gwan Lee, Yeon Sook Kim

## Abstract

Hydrogen bonding magnetic resonance-based cyclic electromagnetic DNA simulation (CEDS) was proposed to play a role in regulating the function of target short oligo-dsDNA in the previous study, as the decagonal CEDS was found to induce the hybridization and conformational changes of target short oligo-dsDNA in a sequence-specific manner. This study applied dodecagonal CEDS on 24 bps oligo-dsDNAs and plasmid DNAs. CEDS can affect 24 bps oligo-dsDNAs intercalated by ethidium bromide (EtBr) or condensed by spermidine by stimulating them to partially remove EtBr or spermidine, respectively, depending on CEDS time. When the multiple cloning site of pBluescript was digested with eight restriction endonucleases (RE) under CEDS separately, all RE digests were enhanced depending on CEDS time compared to the negative and positive controls both in EtBr intercalated electrophoresis and HPLC analysis. *In vitro* RNA transcription from human VEGFA, vWF or elafin cDNA subcloned into pBluescript DNA was increased by CEDS using each promoter sequence depending on CEDS time compared to negative and positive controls. In addition, the productions of green fluorescent protein (GFP) and β-galactosidase were enhanced by CEDS using each promoter sequence from pE-GFP-1 and pBluescript, respectively. In the survival assay using E. coli transfected with pBluescript containing ampicillin resistance gene, the number of surviving E. coli was increased by CEDS using a T3 promoter sequence compared to the control during two days of culture, while it was decreased by CEDS using a T7 promoter, non-specific 12A, 6(TA), or mutation-2 (GG18,19CC) T3 promoter sequence. Therefore, it is suggested that dodecagonal CEDS can target 24 bps oligo-dsDNAs and promoter sequences of plasmid DNAs in a sequence-specific manner and influence their functions.

---

The CEDS is designed to target the base pairs of dsDNA randomly distributed in the buffer solution by the direction of the magnetic field depending on the base pair and the angular momentum of the base pairs per revolution in a decagonal or dodecagonal fashion. In the previous study, decagonal CEDS can target 6-12 bps oligo-dsDNAs in 0.1M NaCl solution that have little tertiary structure. In this study, dodecagonal CEDS was used to target 24 bps oligo-dsDNAs and plasmid DNAs that may have coiled coil and tertiary structures.

## CEDS effect on the binding of ethidium bromide (EtBr) and spermidine in oligo-dsDNAs

### 1) CEDS stimulated oligo-dsDNAs to recover from EtBr intercalation

Since UV260 absorbance is variable depending on the structural changes of dsDNA stiffened by EtBr intercalation in buffer solution (*1*), this study indirectly detected the CEDS-induced conformational changes of oligo-dsDNA in 0.1M NaCl solution with HPLC equipped with UV and fluorescence spectroscopy. The palindromic ss4(3T3A) and pairs of ss6(2C2A) and ss6(2T2G), ss6C6A and ss6T6G were separately dissolved in 0.1M NaCl solution at 100 pmole/μL and hybridized by heating up to 90°C and slowly cooling to room temperature.

The ds4(3T3A) in 0.1M NaCl solution was mixed with EtBr at 1 or 4 μg/mL at room temperature for 10 min, and treated with dodecagonal 4(3T3A)*-CEDS at 20-25 Gauss for 180 min. During the CEDS, 50 μL of ds4(3T3A) solution was obtained, and immediately analyzed by HPLC with a reverse phase silica column and a running buffer of 0.1M NaCl solution at a flow rate of 0.3 mL/min. As ds4(3T3A) was intercalated by 1 and 4 μg/mL EtBr, the peak of ds4(3T3A) decreased up to 93.1% and 75.2% of the untreated control level, respectively, and recovered by 4(3T3A)*-CEDS up to 98.5% and 82.2% until 180 min, respectively (Fig. 1A).

**Fig. 1.**
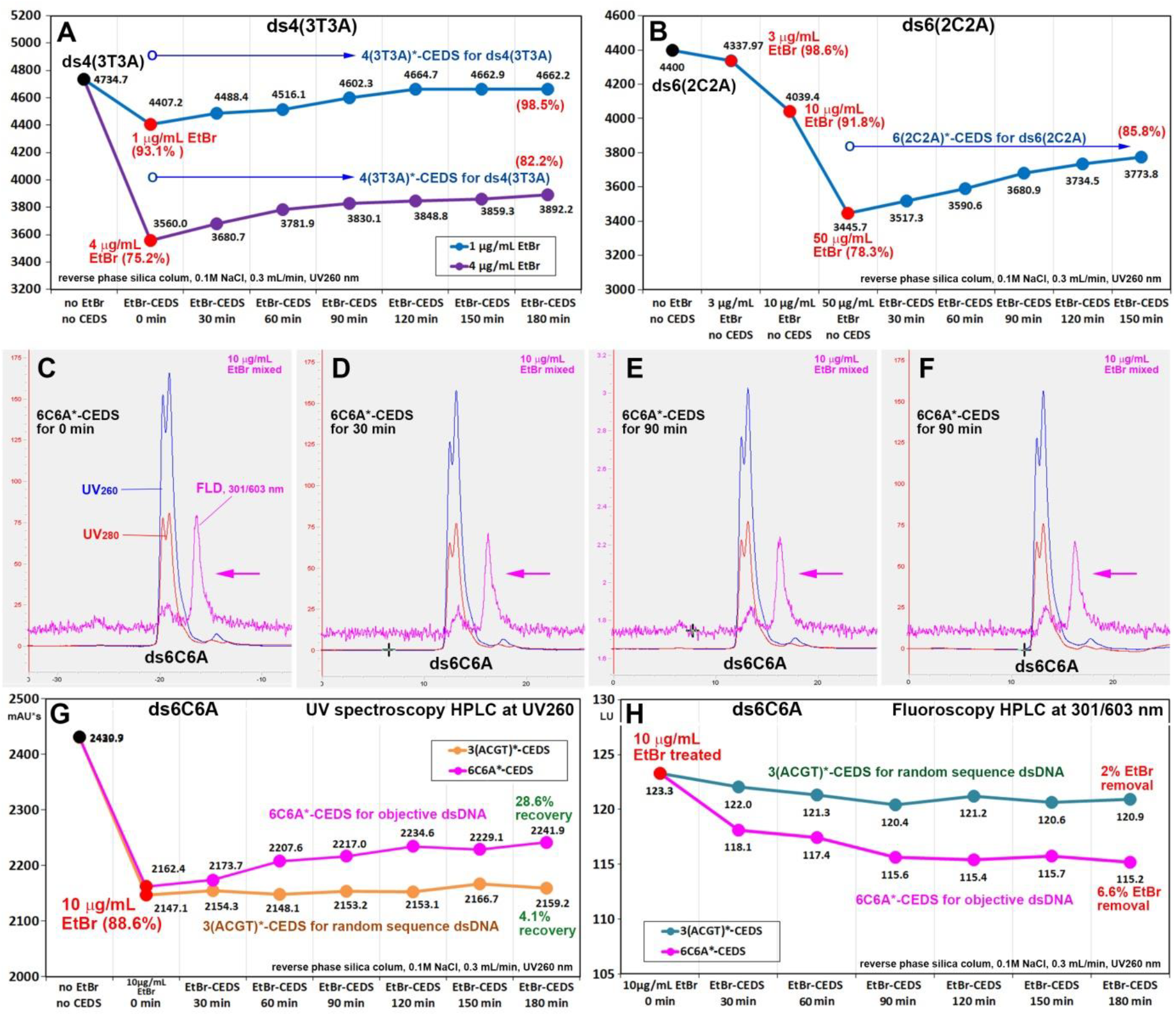
CEDS effect on EtBr intercalation with oligo-dsDNAs in 0.1M NaCl solution through UV and fluorescence HPLC. A: ds4(3T3A) with 1 or 4 μg/mL EtBr. B: ds6(2C2A) with 3, 10, or 50 μg/mL EtBr. C-F: ds6C6A with 10 μg/mL EtBr. G and H: ds6C6A with 10 μg/mL EtBr.

The ds6(2C2A) in 0.1M NaCl solution was mixed with EtBr at 3, 10, and 50 μg/mL by adding standard EtBr solution (1 mg/mL) at room temperature for 10 min separately. And then each sample was treated with dodecagonal 6(2C2A)*-CEDS for 150 min. 50 μL of ds6(2C2A) solution was obtained during CEDS, and immediately analyzed by HPLC as described above. The peak area of ds6(2C2A) decreased up to 98.6%, 91.8%, and 78.3% of untreated control level with increasing concentrations of 3, 10, and 50 μg/mL EtBr, respectively. The maximum decrease induced by 50 μg/mL EtBr was gradually recovered by 6(2C2A)*-CEDS up to 85.8% of untreated control level at 150 min (Fig. 1B).

To determine the EtBr removal from oligo-dsDNA by CEDS, ds6C6A solution was mixed with EtBr at 10 μg/mL for 10 min. and treated with dodecagonal 6C6A*-CEDS or 3(ACGT)*-CEDS at room temperature for 180 min. During the CEDS, 50 μL of ds6C6A solution was obtained and immediately analyzed by HPLC equipped with both UV spectroscopy and fluoroscopy using the method described above. As a result, the ds6C6A solution with 10 μg/mL EtBr resulted a decrease in peak area up to 88.6% of the untreated control level, was reversed by 6C6A*-CEDS, up to 28.6% at 180 min, and was weakly reversed by random sequence 3(ACGT)*-CEDS, up to 4.3% (Fig. 1G). In HPLC fluoroscopy analysis for EtBr fluorescence at 301 nm excitation and 603 nm emission, EtBr fluorescence was decreased by 6C6A*-CEDS up to 6.6% of untreated control level at 180 min, while slightly decreased by 3(ACGT)*-CEDS up to 2% (Fig. 1 C-F,H). The results indicate the EtBr bound to oligo-dsDNA was partially removed by CEDS in a sequence specific manner.

### 2) CEDS stimulated oligo-dsDNAs to recover from the spermidine condensation

Spermidine readily binds to dsDNA, causing condensation and conformational changes (*2*), which can be detected by UV260 spectroscopy. The ds3(4C4A) in 0.1M NaCl solution at 100 pmole/μL was prepared as described above, and mixed with spermidine at varying concentrations of 1-100 mM separately. Each sample was treated with dodecagonal 3(4C4A)*-CEDS for 30 min and then analyzed by HPLC using the method described above. The samples mixed with spermidine at various concentrations showed decreases in peak area depending on the concentration of spermidine, resulting in a steep downward slope up to 77% of untreated control level at 100mM spermidine (Fig. 2 A-C). Whereas when the samples mixed with spermidine at various concentrations were separately treated with 3(4C4A)*-CEDS, their peak area showed a relatively slow decrease with a gentle downward slope up to 86. 2% of untreated control level compared to untreated control (Fig. 2C).

**Fig. 2.**
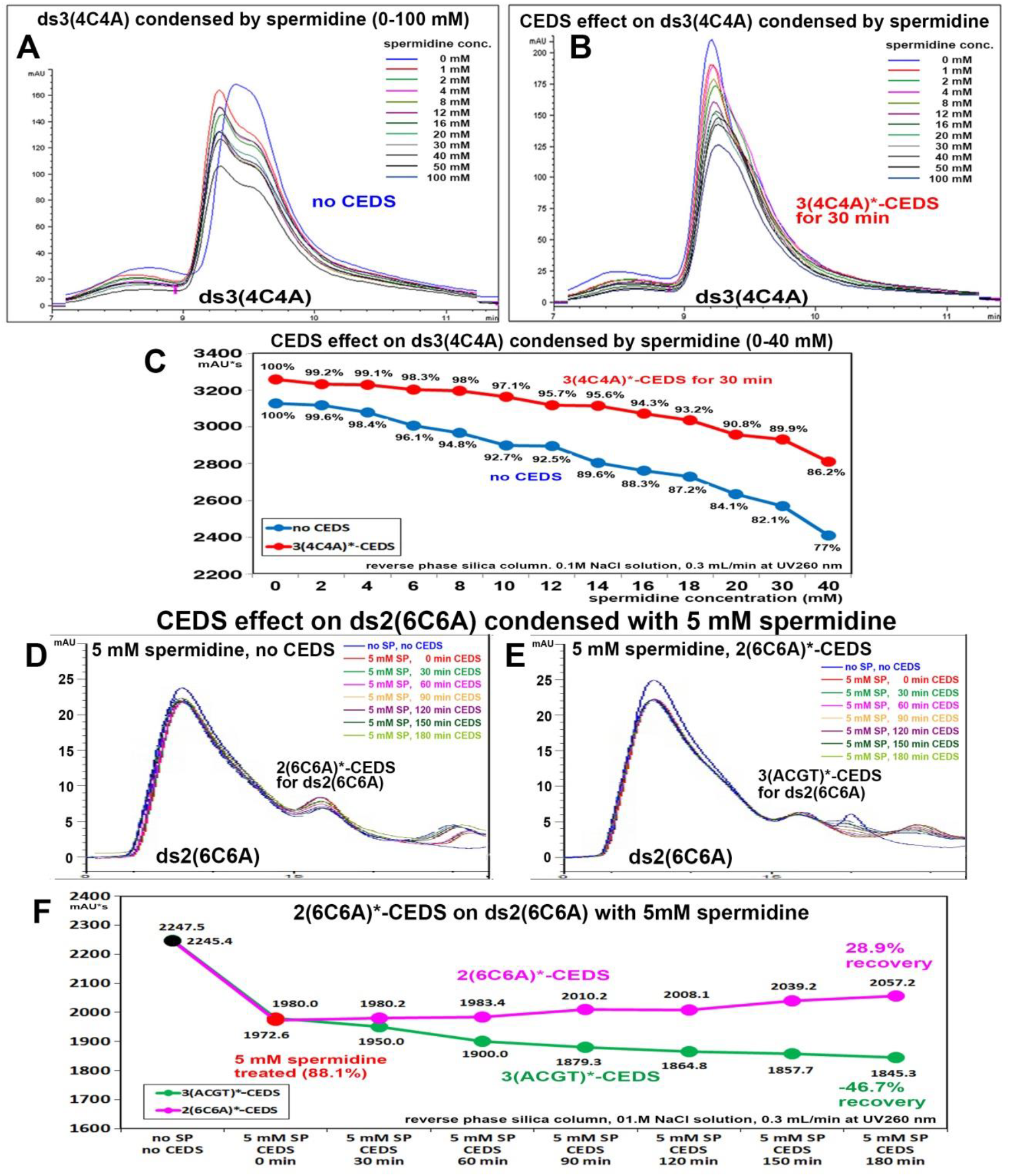
CEDS effect on the spermidine-induced DNA condensation of oligo-dsDNA through HPLC analysis. Spermidine-induced oligo-dsDNA condensation was recovered by CEDS. A-C: ds3(4C4A) with 0-100 mM spermidine. D-F: ds2(6C6A) with 5 mM spermidine.

To know the CEDS effect on spermidine-induced oligo-dsDNA condensation depending on CEDS time, ds2(6C6A) in 0.1M NaCl solution was mixed with spermidine at 5 mM for 10 min, and treated with dodecagonal 2(6C6A)*-CEDS or 3(ACGT)*-CEDS at room temperature for 180 min. During the CEDS, 50 μL of ds2(6C6A) sample was obtained, and immediately analyzed by HPLC using the method described above. The ds2(6C6A) showed a marked reduction of peak area up to 88.1% of untreated control level by mixing with spermidine at 5 mM concentration, and the reduced peak area was gradually recovered by 2(6C6A)*-CEDS up to 28.9% of the spermidine-induced decrease at 180 min, whereas it was rather decreased by 3(ACGT)*-CEDS up to 46.7% (Fig. 2 D-F).

The data indicate that spermidine significantly condensed ds2(6C6A), and this condensation was reversed by 2(6C6A)*-CEDS, but not by 3(ACGT)*-CEDS. Therefore, it is proposed that CEDS can renature ds2(6C6A) to partially recover from the spermidine-induced deformation of DNA in a sequence-specific manner.

### CEDS enhanced the restriction endonuclease (RE) digestion of plasmid DNAs

The multiple cloning site of pBluescript SK(-) was separately digested with BamHI, EcoRI, KpnI, NotI, PstI, SacI, XhaI and XhoI enzymes (New England Biolabs, USA) (*3*) under decagonal GGATCC*-CEDS, GAATTC*-CEDS, GGTACC*-CEDS, GCGGCCGC*-CEDS, CTGCAG*-CEDS, GAGCTC*-CEDS, TCTAGA*-CEDS and CTCGAG*-CEDS, respectively, at 37°C for 30 min. The DNA product underwent electrophoresis on 1% agarose gel, stained with EtBr, and was detected under UV illumination. The negative and positive controls were performed simultaneously without CEDS and with decagonal 2(ACGT)*-CEDS, respectively.

All RE digestions under CEDS using a target sequence showed the stronger DNA bands compared to the negative and positive controls, although there was a slight difference in the degree of band density (Fig. 3 A,C-I). However, when the DNA products of BamHI and XhoI digestion under GGATCC*-CEDS and CTCGAG*-CEDS, respectively, were examined by pre-electrophoresis EtBr staining, they clearly showed stronger DNA bands than the negative and positive controls (Fig. 3 B,J). The data indicate that the CEDS can enhance RE digestion in a sequence-specific manner.

**Fig. 3.**
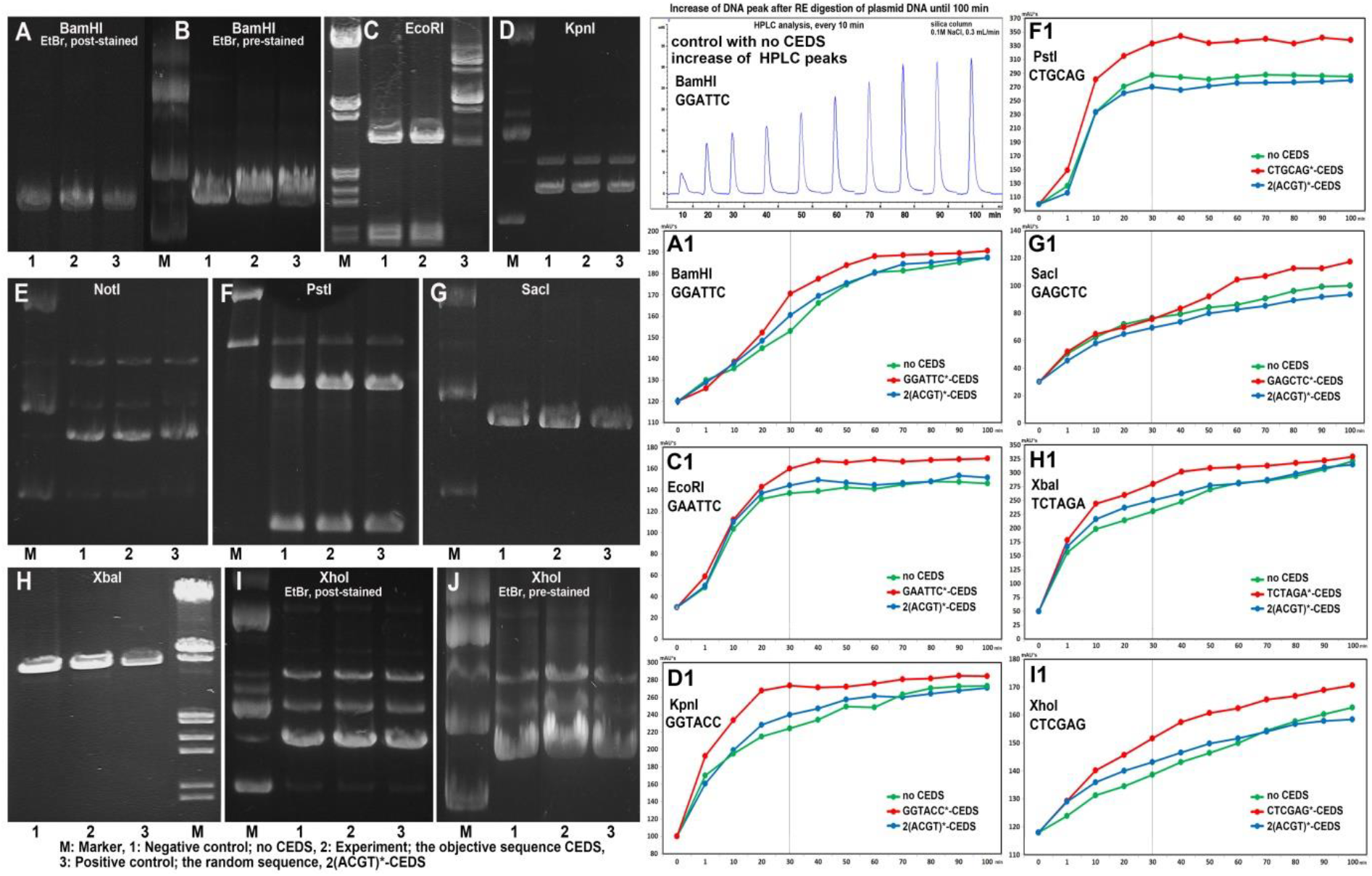
Electrophoresis (A-J) and HPLC analysis (A1-I1) for CEDS effect on different RE digestions of pBluescript SK(-). RE digestion was increased by CEDS compared to untreated control and 2(ACGT)*-CEDS depending on CEDS time.

To evaluate CEDS-induced RE digestion depending on CEDS time, each 30 μg of plasmid DNA was mixed with 300 μL of RE digestion mixture, and separately digested with 30 unit of BamHI, EcoRI, KpnI, PstI, SacI, XhaI, and XhoI, under CEDS using each sequence in the incubator at 37°C for 100 min. Additionally, RE digestion without CEDS and with random sequence (2(ACGT))*-CEDS were performed simultaneously. During the RE digestion under CEDS, each 20 μL of DNA sample was obtained, and analyzed by HPLC using a reverse phase silica column, 0.1M NaCl solution, 0.5 mL/min, at UV260.

As a result, RE digestion of plasmid DNA with BamHI, EcoRI, KpnI, PstI, SacI, XbaI, or XhoI enzyme was rapidly increased by CEDS during 1-30 min of CEDS time and plateaued at 60 min compared to the negative and positive controls (Fig. 3 A1, C1, D1, F1-I1). The data show RE digestion of plasmid DNA was enhanced by CEDS in the early stage until 30 min, resulting in significant higher RE digestion by CEDS in the late stage until 100 min than by no CEDS and 2(ACGT)*-CEDS.

### CEDS increased *in vitro* RNA transcription from plasmid DNA inserted with human VEGFA, vWF, or elafin cDNAs

*In vitro* RNA transcription was performed using linear plasmid DNAs (pBluescript SK(-)) inserted with human VEGFA, vWF, or elafin cDNAs (500-600 bps) using *in vitro* transcription reaction kit (Enzynomics, Korea). Each sample mixture was treated with dodecagonal T7 promoter-CEDS (TAATACGACTCACTATAGGG-CEDS), SP6 promoter-CEDS (ATTTAGGTGACACTATA-CEDS), or T3 promoter-CEDS (AATTAACCCTCACTAAAGGG-CEDS) at 20-25 Gauss, 37°C for 20 and 40 min. After the experiment, the mixtures were treated with DNase I to degrade the template DNAs. The RNA products were then immediately electrophoresed using 1.5% formaldehyde agarose gel (20 mM MOPS, 6% formaldehyde) and DEPC-based buffer, stained with EtBr, and detected under UV illumination.

The study found that *in vitro* RNA transcription of VEGFA, vWF, and elafin under T7 promoter-CEDS, SP6 promoter-CEDS, or T3 promoter-CEDS, respectively, increased up to 8.9%, 12.9%, and 33.4% at 20 min and up to 6.2%, 2.4%, and 21.8% at 40 min, respectively, compared to untreated control (Fig. 4 A-F). Particularly, forward SP6 promoter-CEDS for 20 min increased *in vitro* RNA transcription of vWF up to 21.1% more than untreated control, whereas reverse SP6 promoter-CEDS increased only a 9% increase (Fig. 4 G,H).

**Fig. 4.**
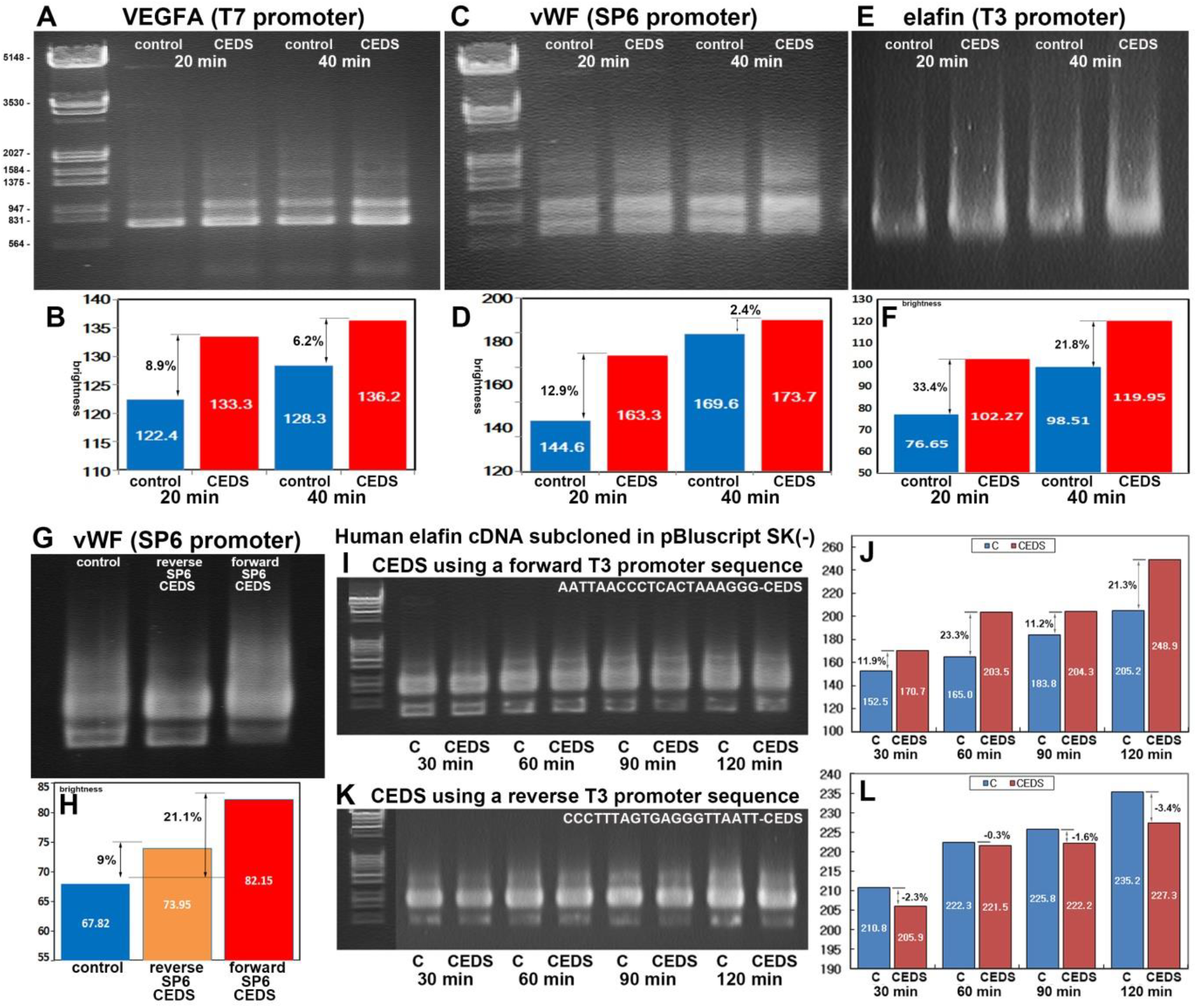
*In vitro* RNA transcription from plasmid DNA inserted with human VEGFA (A and B), vWF (C,D,G,H), elafin (E,F,I-L) cDNAs under T7 promoter-CEDS, SP6 promoter-CEDS, and T3 promoter-CEDS, respectively.

To identify the sequential increase of *in vitro* RNA transcription depending on CEDS time, 10 μg of pBluescript DNA containing human elafin cDNA was dissolved in 100 μL of reaction mixture, and treated with dodecagonal forward T3 promoter sequence-CEDS (AATTAACCCTCACTAAAGGG-CEDS) or reverse T3 promoter sequence-CEDS (CCCTTTAGTGAGGGTTAATT-CEDS) in an incubator at 37°C for 120 min. During the incubation under CEDS, 10 μL of the reaction mixture was taken, and then immediately electrophoresed using the procedure described above.

Forward T3 promoter-CEDS increased *in vitro* RNA transcription of elafin by 11.9%, 23.3%, 11.2%, and 21.3% at 30, 60, 90, and 120 min, respectively, whereas reverse T3 promoter-CEDS decreased by 2.3%, 0.3%, 1.6%, and 3.4%, respectively (Fig. 4 I-L). The data indicate that forward T3 promoter-CEDS consistently enhanced RNA transcription for 120 min, while reverse T3 promoter-CEDS slightly inhibited RNA transcription.

### CEDS increased the reporter protein production of plasmid DNA vectors

Before the CEDS targeting each promoter sequence in plasmid DNA, we performed the conformational assay for oligo-ssDNA and oligo-dsDNA treated with dodecagonal CEDS in 0.1M NaCl solution, and found that both T3 promoter oligo-ssDNA and oligo-dsDNA are more responsive to forward T3 promoter-CEDS by increasing up to 14.9% and decreasing up to 14.5%, respectively, compared to random sequence 3(ACGT)-CEDS (Supplementary Text, Fig. S1). The results indicate that forward T3 promoter CEDS affects oligo-ssDNA and oligo-dsDNA in reverse by extending and condensing their conformation, respectively, in contrast to 3(ACGT)-CEDS.

### CEDS-induced production of green fluorescent protein (GFP) from pE-GFP-1

E. coli were transfected with pE-GFP-1 vector (Clontech, USA) containing a GFP gene at the 5’ flanking end of the T3 promoter, and cultivated in Luria Bertani (LB) medium supplemented with kanamycin (50 μg/mL). The E. coli culture was prepared at a standard concentration, 0.5 at OD600. The standard E. coli culture was incubated under dodecagonal T3 promoter-CEDS at 10 Gauss, 37°C for 3 and 5 hours. The positive and negative controls were also incubated under dodecagonal random sequence-CEDS (6(ACGT)-CEDS) and no CEDS, respectively. After CEDS treatment, the supernatant was obtained by centrifugation at 1000g for 10 min, and the production of GFP was measured by a spectrofluorometer (Jasco, Japan) with 488 nm excitation and 507 nm emission.

The culture products were also compared on nitrocellulose membrane by observing under UV light (Fig. S2), however, the spectrofluorometer analysis showed that GFP production under T3 promoter-CEDS significantly increased by approximately 70.6% and 66.7% at 3 and 5 hours of culture, respectively, compared to the negative control performed without CEDS, while GFP production under 6(ACGT)-CEDS showed an increase of only 33.3% and 6.1%, respectively (Fig. 5 A). The data suggest that T3 promoter-CEDS may activate T3 promoter and result the overproduction of GFP compared to the negative and positive controls.

**Fig. 5.**
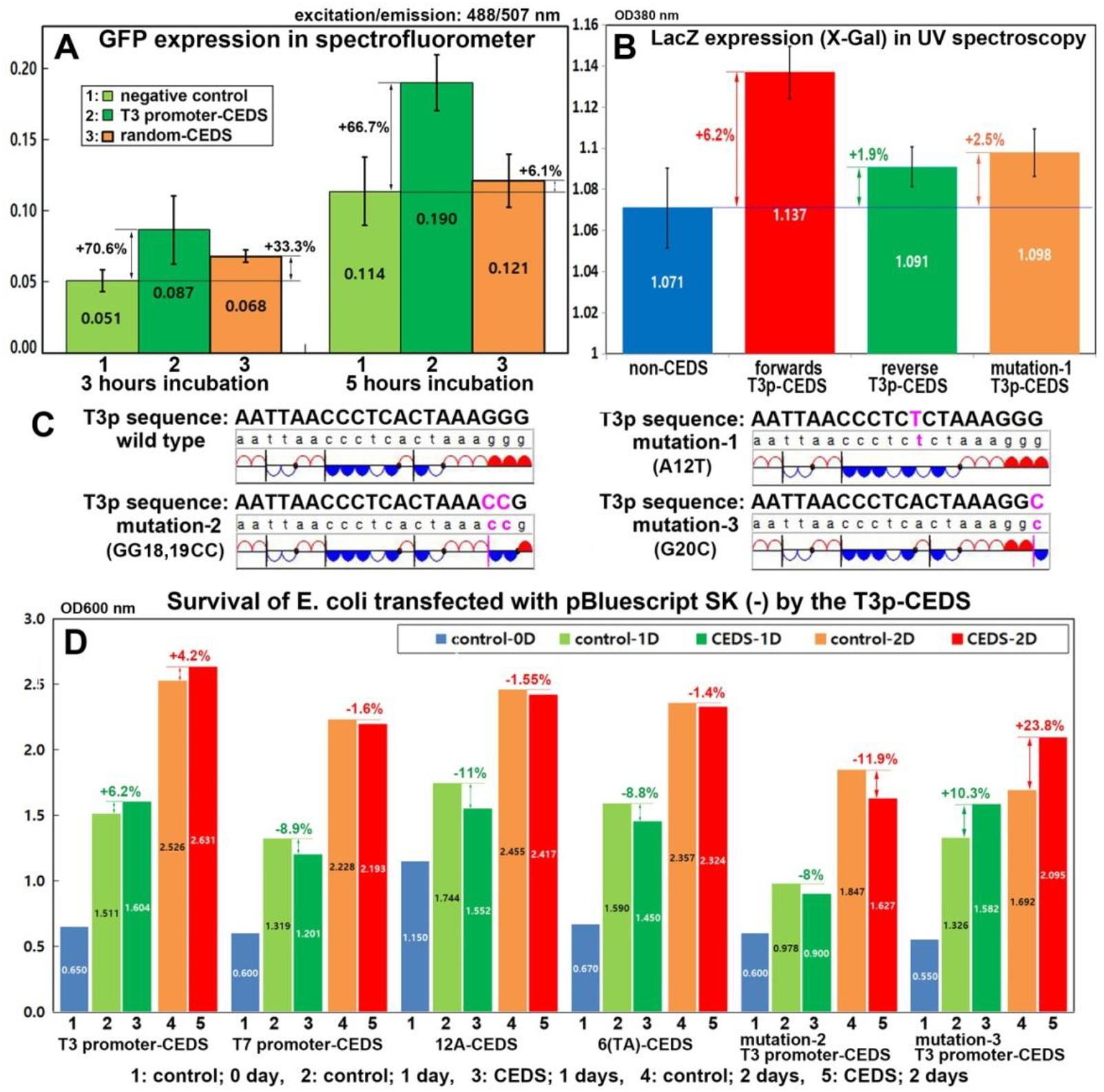
CEDS effect on the expression of green fluorescent protein (GFP) and β-galactosidase from pBluescript SK(-) and pE-GFP-1 vector, respectively (A,B). Spectroscopy analysis for E. coli survival in ampicillin-supplemented LB media depending on dodecagonal T3 promoter-CEDS, T7 promoter-CEDS, 12A-CEDS, 6(TA)-CEDS, mutation-2 T3 promoter-CEDS or mutation-3 promoter-CEDS (C,D).

### CEDS-induced production of β-galactosidase from pBluescript vector and cell survival

The pBluescript SK(-) vector from Stratagene (USA) contains a LacZ gene under the 5’ and 3’ flanking T3 and T7 promoters, respectively. The T3 promoter encodes the LacZ and ampicillin-resistance genes in the forward direction, while the reverse direction does not (*4*). The standard E. coli culture mixture was generated by introducing the vector into E. coli and cultivating it in LB media supplemented with 100 μg/mL ampicillin, 0.5 at OD600. 1.5 mL of the standard E. coli culture mix in a 15 mL tube was treated with dodecagonal forward T3 promoter-CEDS, reverse T3 promoter-CEDS, or mutation-1 (A12T) T3 promoter-CEDS at room temperature for 12 hours. A weak magnetic field, 2-3 Gauss, was utilized for prolonged CEDS treatment on cells. The β-galactosidase production was detected by mixing with 1 mM IPTG and 10 mM X-Gal, and the resulting blue color was immediately measured by spectrophotometer at 380 nm. The X-Gal reaction of culture products can be observed visually (Fig. S3).

Forward T3 promoter-CEDS was observed to increase β-galactosidase production in cells by up to 6.2% compared to untreated control, while reverse T3 promoter-CEDS demonstrated a slight increase of up to 1.9% in cells. In contrast, the mutation-1 (A12T) T3 promoter-CEDS exhibited a slight increase in β-galactosidase production up to 2.5% (Fig. 5B). The data indicates that forward T3 promoter-CEDS enhances the transcription of LacZ gene in a sequence-specific manner from pBluescript vector in cells, in comparison to reverse T3 promoter-CEDS or mutation-1 (A12T) T3 promoter-CEDS.

T3 promoter activation of pBluescript is critical for the survival of E. coli in ampicillin-supplemented media. To evaluate cell survival, 1.5 mL culture of E. coli transfected with pBluescript SK(-) vector, 0.5 at OD600, was prepared in a 15 mL tube as described previously. The culture was treated with T3 promoter-CEDS, T7 promoter-CEDS, 12A-CEDS, 6(TA)-CEDS, along with a mutation-2 (GG18,19CC) T3 promoter-CEDS, or mutation-3 (G20C) T3 promoter-CEDS (Fig. 5C). The CEDS treatment was carried out using low magnetic field, 2-3 Gauss, at room temperature for 1 or 2 days. After treatment, the culture was immediately examined for surviving cells by measuring OD600 with spectrophotometer.

As a result, T3 promoter-CEDS increased the OD600 absorbance up to 6.2% after 1 day and 4.2% after 2 days compared to untreated control. In contrast, T7 promoter-CEDS, 12A-CEDS, or 6(TA)-CEDS decreased the OD600 absorbance by 8.9%, 11%, and 8.8% after 1 day and 1.6%, 1.55%, and 1.4% after 2 days, respectively. On the other hand, mutation-2 (GG18,19CC) T3 promoter-CEDS decreased the OD600 absorbance up to 8% after 1 day and 11.9% after 2 days, while mutation-3 (G20C) T3 promoter-CEDS increased the OD600 absorbance up to 10.3% after 1 day and 23.8% after 2 days (Fig. 5D).

The data indicate that T3 promoter-CEDS enhanced the transcription of ampicillin resistance gene, allowing bacteria to survive in ampicillin-supplemented media in contrast to T7 promoter-CEDS, nonspecific sequences 12A-CEDS, and 6(TA)-CEDS. Moreover, CEDS using a mutation-2 (GG18,19CC) sequence, which has been replaced with a Pyu dsDNA segment at the 3’ end of T3 promoter, has more negative impact than CEDS using a mutation-3 (G20C) T3 promoter sequence, which has been mutated only at the end of T3 promoter.

### Ampicillin resistance protein production by CEDS using a selected promoter sequence

For the survival assay of E. coli transfected with pBluescript vector, dodecagonal CEDS can use a sequence directly selected from promoter region of ampicillin resistance gene in pBluescript through the analysis of DNA base pair polarity program (Fig. 6 A,D). Simply, a forward selected-sequence (dsATTCAAATATGTATCCGCTCATGA) consisting of 7 Pyu dsDNA segments with even base pair polarities was applied to dodecagonal CEDS generating 2-3 Gauss at room temperature for 18 hours on 1.5 mL standard of E. coli culture mix as described above. On the other hand, dodecagonal CEDS using a reverse selected-sequence (dsTCATGAGCGGATACATATTTGAAT) was simultaneously performed. After treatment, the number of surviving cells was determined by analyzing the culture products using spectrophotometer at OD600.

**Fig. 6.**
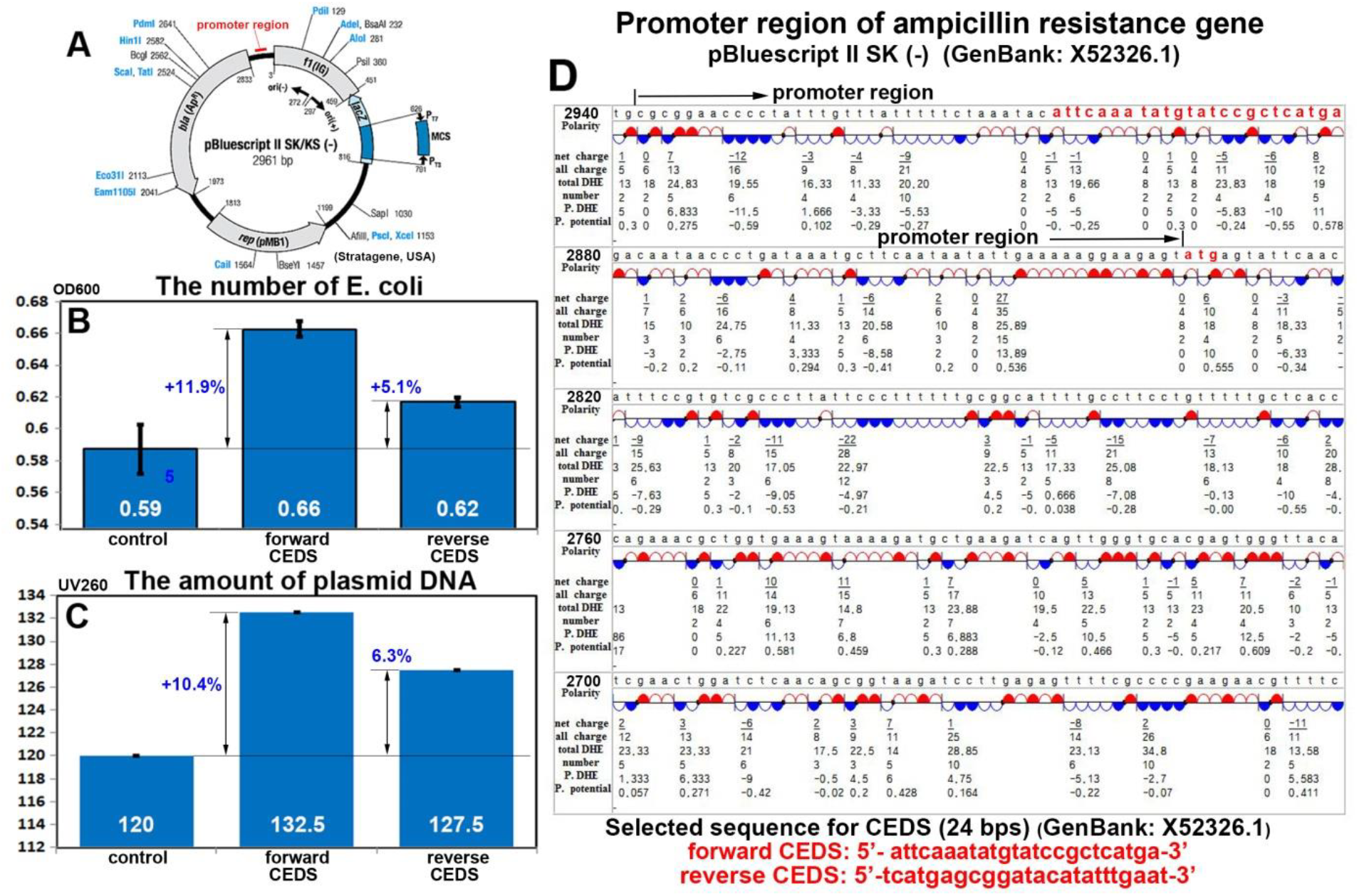
The survival of E. coli transfected with pBluescript II SK(-) vector in LB media supplemented with ampicillin increased by CEDS using a 24 bp sequence, consisting of 7 Pyu dsDNA segments with even polarities, selected from the promoter region of ampicillin resistance gene.

The CEDS using a forward selected-sequence (dsATTCAAATATGTATCCGCTCATGA) increased the number of E. coli by 11.9% after 18 hours compared to the untreated controls. In contrast, CEDS using a reverse selected-sequence (dsTCATGAGCGGATACATATTTGAAT) increased the cell count by only 5.1% (Fig. 6 B).

On the other hand, the plasmid DNAs were purified from the culture products and examined by spectrophotometer at UV260 to determine the amount of plasmid DNA by the CEDS. The CEDS using the forward selected-sequence above increased the amount of plasmid DNA by 10.4% after 18 hours compared to untreated controls. In contrast, the CEDS using the reverse selected-sequence increased the plasmid DNA amount by only 6.3% (Fig. 6 B,C).

The data show that CEDS using uniformly polarized multiple Pyu dsDNA segments located at the track of the transcription complex can enhance transcription of the target gene similarly to CEDS using its exact promoter sequence, such as the T3 promoter. Therefore, it is suggested that the application of CEDS for objective gene regulation may be much more versatile than we expected.

## Discussion

This study applied dodecagonal CEDS to 24 bps oligo-dsDNAs and plasmid DNAs, and found that CEDS can influence target oligo-dsDNA to recover from EtBr- and spermidine-induced conformational change, affect plasmid DNA by enhancing RE digestion and RNA transcription *in vitro*, and more, it can increase the production of GFP and β-galactosidase, and ampicillin resistance protein from plasmid DNAs.

However, the results of CEDS targeting long oligo-dsDNAs and plasmid DNAs in salt buffer solution for EtBr and spermidine binding assay, RE digestion and *in vitro* RNA transcription assay were relatively proportional to the function of CEDS, while the results of CEDS targeting plasmid vectors in prokaryote cells, E. coli, for the production assays of GFP and β-galactosidase and ampicillin resistance protein appears to be irregular and variable. Therefore, further investigation is required to apply CEDS to target dsDNAs in mammalian cells in the following studies.

## Acknowledgments

We would like to express our gratitude to the late Professor Je Geun Chi and the late Dr. Soo Il Chung, who contributed to this research in part.

