## Supplement Fig S1-S3 for "Cyclic Electromagnetic DNA Simulation (CEDS) influences the functions of Long Oligo-dsDNAs and Plasmid DNAs"

### The PDF file includes:

Materials and Methods
Supplementary Text
Figs. S1 to S3
References

### Materials and Methods

#### *CEDS effect on single- and double-strand T3 promoter DNA*

The forward T3 promoter oligo-ssDNA (AATTAACCCTCACTAAAGGG) ssDNA and T3 promoter oligo-dsDNA were separately prepared in 0.1M NaCl solution at 100 pmole/μL by heating to 90℃ and slowly cooling to room temperature. The samples were then treated with dodecagonal CEDS using a forward T3 promoter sequence or a random sequence, 3(ACGT), for 30 min at room temperature. The resulting DNA products were immediately examined with spectrophotometer (Optizen, Korea) at UV260 (Fig. S1).

#### *CEDS-induced GFP expression detected on nitrocellulose membrane*

E. coli cells were transfected with the pE-GFP-1 vector (Clontech, USA) and cultivated in Luria Bertani (LB) medium supplemented with kanamycin (50 μg/mL). The pE-GFP-1 vector contains a green fluorescent protein (GFP) gene, inserted at the 5' flanking end of the T3 promoter. The E. coli cell culture was prepared at 0.5 OD600, and incubated under dodecagonal T3 promoter sequence-CEDS at 20-25 Gauss, 37℃ for 30 min. The control groups were also incubated with no CEDS. After CEDS treatment, the E. coli culture was incubated in a shaking incubator at 37℃ for 1 hour. The supernatant was obtained after centrifugation at 1000g, and 50 μL of culture product was placed on nitrocellulose membrane and observed under UV light (Fig. S2).

***CEDS-induced β-galactosidase production of pBluescript SK(-) vector***

The pBluescript II SK(-) vector from Stratagene (USA) contains a LacZ gene under the 5' and 3' flanking T3 and T7 promoters, respectively. The β-galactosidase assay is a popular gene regulation experiment that employs a LacZ gene-containing vector, making it both unique and simple to conduct. This study assesses the expression of β-galactosidase from the LacZ gene of the pBluescript vector when treating dodecagonal CEDS using a T3 or T7 promoter sequence.

The E. coli culture transfected with pBluescript SK (-) vector was prepared as described above. Then, it was treated with CEDS using a forward T3 promoter, mutated T3 promoter (GG18,19CC), and poly-A 22A sequence at room temperature for 2 days. The treated culture was immediately examined by spectrophotometer at UV 380 nm for X-Gal reaction. A weak magnetic field (3-5 Gauss) was utilized for prolonged exposure on cells during this process. To perform the β-galactosidase assay, mix the culture product with 1 mM IPTG and 10 mM X-Gal. Detect the X-Gal reaction at 380 nm using a UV-spectrophotometer and subject it to statistical analysis.

### Supplementary Text

#### *CEDS effect on single- and double-strand T3 promoter DNA*

Single-strand T3 promoter DNA (T3 promoter oligo-ssDNA) and double-strand T3 promoter DNA (T3 promoter oligo-dsDNA) in 0.1M NaCl solution differently responded to T3 promoter-CEDS. T3 promoter oligo-ssDNA showed a significant increase in UV260 absorption by up to 14.9% in forward T3 promoter-CEDS compared to untreated control, while a weak increase by up to 4.8% by random sequence 3(ACGT)-CEDS (Fig. S1A). Whereas T3 promoter oligo-dsDNA showed a significant decrease in UV260 absorption by up to 14.5% by the forward T3 promoter-CEDS compared to untreated control while only a weak increase by up to 0.9% by 3(ACGT)-CEDS (Fig. S1B).

The results indicate that both T3 promoter oligo-ssDNA and oligo-dsDNA in 0.1M NaCl solution are more responsive to forward T3 promoter-CEDS by increasing up to 14.9% and decreasing up to 14.5%, respectively, than to 3(ACGT)-CEDS.

#### *CEDS-induced GFP expression detected on nitrocellulose membrane*

The experimental group subjected to CEDS treatment using a T3 promoter sequence to target GFP gene in pE-GFP-1 vector consistently generated more GFP than the positive and negative control groups for up to 5 hours during the experiment. When the culture of E. coli transfected with pE-GFP-1 vector was treated with CEDS using a T3 promoter sequence at low magnetic field, 10 Gauss, for 3 and 5 hours, the culture products showed stronger GFP fluorescence on nitrocellulose membrane by 20% and 16.7%, respectively, than untreated control (Fig. S2A-C).

***CEDS-induced β-galactosidase production of pBluescript SK(-) vector***

In the culture of E. coli transfected with pBluescript SK(-) containing β-galactosidase gene (LacZ gene) for 2 days, forward T3 promoter-CEDS increased UV380 absorbance by 5.66% in 1 day and 9.74% in 2 days compared to untreated control, while mutated promoter (GG18,19CC)-CEDS decreased UV380 absorbance by 7.93% in 1 day and 6.87% in 2 days. On the other hand, the nonspecific poly-A sequence, 22A-CEDS also decreased UV380 absorbance by 18.08% in 1 day and 1.55% in 2 days.

This simple experiment of E. coli culture shows that the T3 promoter-CEDS can target T3 promoter sequence in a sequence specific manner in contrast the mutated T3 promoter-CEDS and 22A-CEDS (Fig. S3).

***Discussion***

Before this study, we investigated the magnetic effect on the hydrogen bonding of water, and found characteristic physicochemical properties of magnetically treated water (MTW). The MTW was produced by unidirectional pulsating magnetic field, 800 Gauss, 9 Hertz for 1 hour, and showed the increase of T2 relaxation time in NMR spectroscopy (*1*), the decrease of electrical conductivity and surface tension in degassed state (*1*), the increase of IR absorption at 2400-1900 $cm^{-1}$ (peak at 2115 $cm^{-1}$) (*1*), the increased solubility speed of glycine ($NH_2CH_2COOH$), boric acid ($H_3BO_3$), and $MgSO_4$ but the decreased solubility speed of urea ($CH_4N_2O$), sodium citrate ($HOC(CO_2Na)$-$(CH_2CO_2Na)_2$-$2H_2O$) and $(NH_4)_2SO_4$(*2*), the increase of gypsum crystallization (*3*), the increased precipitation $BaSO_4$, $BaCO_3$, and $CaCO_3$(*3*), the increase of hydration hardening speed of gypsum plaster (*3*), the decrease in critical micelle concentrations (CMC) of sodium dodecyl sulfate (SDS), cetyltrimethylammonium bromide (CTAB), and Pluronic F-68 (*4*), the increase of bony decalcification (*5*), the increase of membrane permeability, and the decrease of free radical activity (*4*).

In the previous studies, we also examined the effect of pulsed unipolar magnetic fields on cells and animals in relation to bio-hazards of alternating magnetic fields. The results showed that the pulsed unipolar magnetic field, with strength of 63-225 Gauss and a frequency of 120 Hertz, had almost no impact on the culture of human osteogenic sarcoma (HOS) cells until 6 hours after magnetic exposure. After this time, apoptotic cell death was observed (*6*). The impact of extremely low frequency electromagnetic field (ELF-EMF) on cells can vary depending on the strength, frequency, and duration of exposure. In a study on mice, exposure to pulsed unipolar ELF-EMF at 730-960 Gauss resulted in rare changes in testicular cells up to 1 hour after exposure, after which the cells gradually underwent apoptotic cell death. Additionally, exposure to 7 Hertz ELF-EMF resulted in more apoptosis of spermatocytes compared to exposure to 1, 20, 40, and 80 Hertz ELF-EMF(*7*). After 2 hours of magnetic exposure with a pulsed unipolar magnetic field of 0.2-0.3T and 60 Hertz, there was almost no expression of amyloid precursor protein (APP) in the mouse brain. However, after 4 hours of magnetic exposure, there was an increase in APP expression (*8, 9*).

The results of the *in vitro* and *in vivo* experiments suggest that the damages caused by the pulsed unipolar magnetic field on cells and animals are due to the occurrence of free radicals resulting from the alternating magnetic field. In fact, the pulsed unipolar magnetic field can induce free radicals that can be used for bony decalcification in histological procedures (*5*). However, the cells and animals remained healthy and exhibited normal features in gross and histological observations for up to 1-2 hours after magnetic exposure (*6-8*). Therefore, it is recommended to limit the strength, frequency, and exposure time of the magnetic field for CEDS in biological applications to prevent potential bio-hazards in cells. In this study, we utilized a pulsed unipolar magnetic field with a frequency of 100 Hertz, strength of less than 30 Gauss, and duration of less than 30 min, as a standard procedure in CEDS.

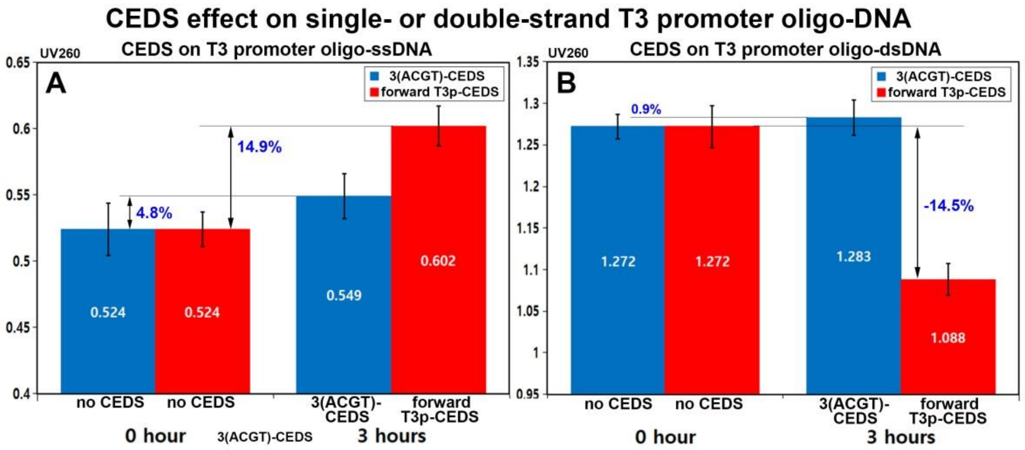


**Fig. S1.** The conformational changes of single or double-strand T3 promoter oligo-DNA by CEDS. T3 promoter oligo-ssDNA (A) and oligo-dsDNA (B) treated with forward T3 promoter-CEDS and random sequence 3(ACGT)-CEDS, respectively.

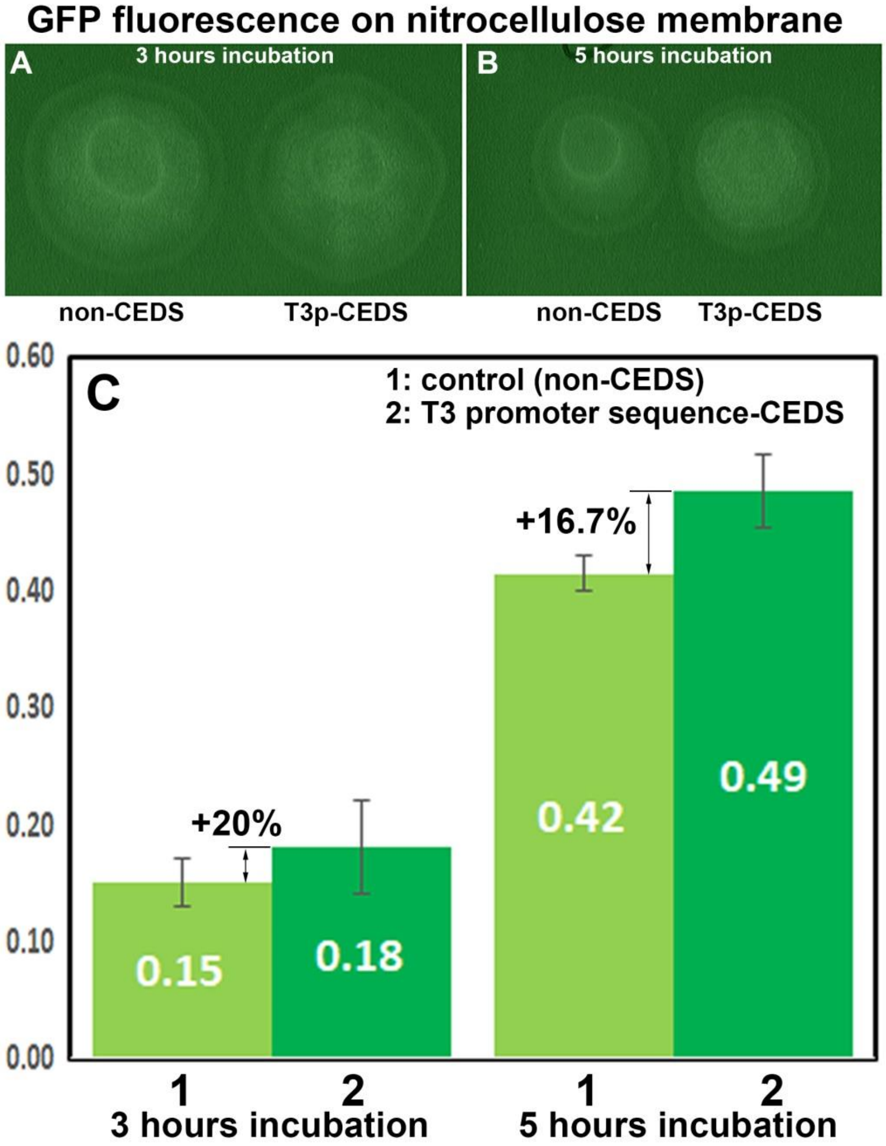


**Fig. S2.** E. coli culture for green fluorescent protein (GFP) expression under CEDS using a T3 promoter sequence. A-C: The comparison between T3 promoter sequence-CEDS and non-CEDS on the culture of pE-GFP-1 vector-transfected E. coli on nitrocellulose membrane (A,B). The densitometer data of A and B were plotted into a graph (C).

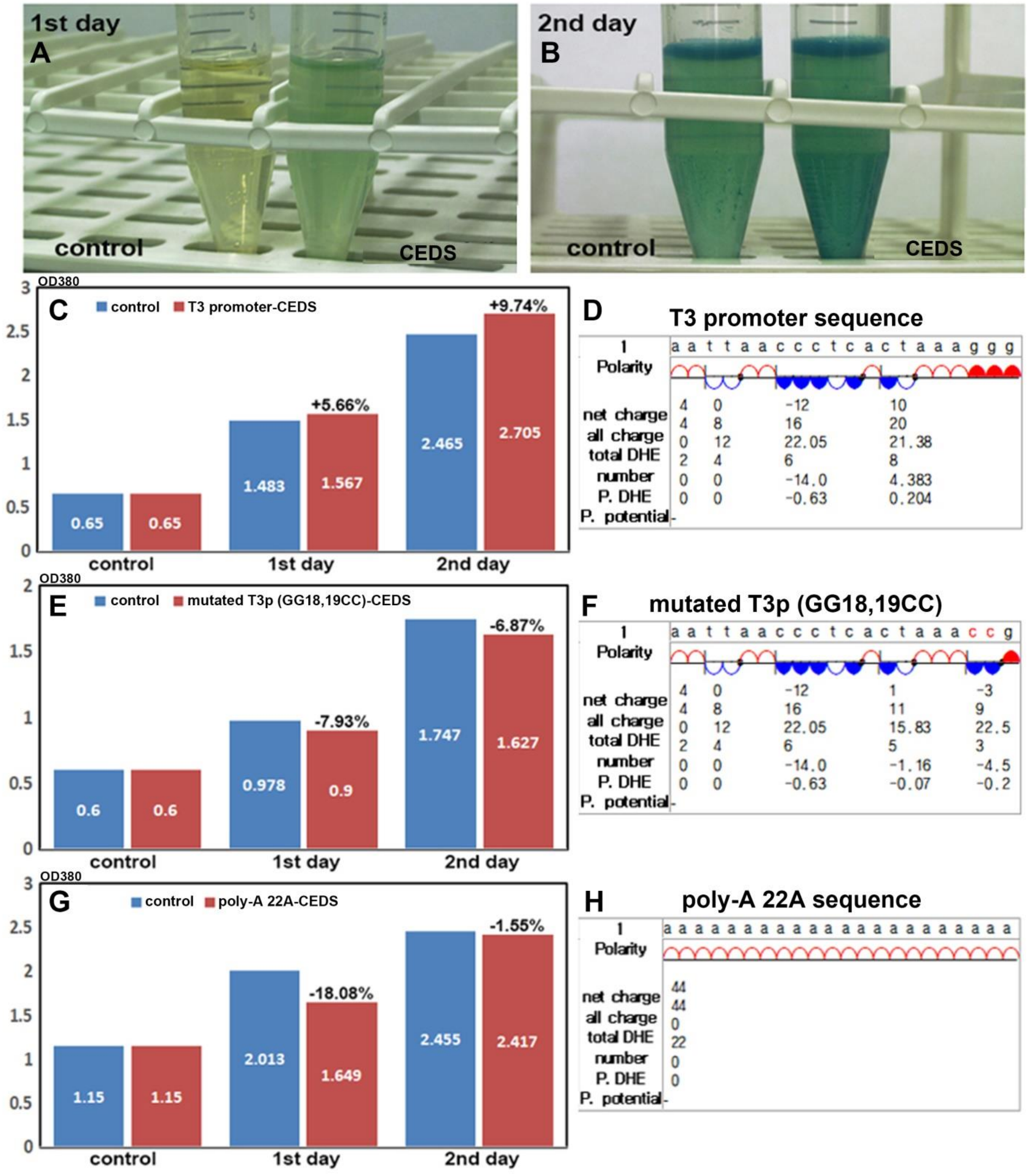


**Fig. S3.** The production of β-galactosidase from E. coli transfected with pBluescript II SK(-) was variable depending on CEDS using a T3 promoter, mutated T3 (GG18,19CC), or poly-A 22A sequence (D,F,H). The X-Gal reaction was visually observed (A,B) and detected by spectrophotometer at OD400 (C,E,G).
